# Plasma oxytocin measured by LC–MS/MS varies with life stage, sex, and obesity in mice

**DOI:** 10.64898/2026.06.25.734250

**Authors:** Georgia Colleluori, Chiara Galli, Simone Moretti, Stefano Di Bona, Ilenia Severi, Jessica Perugini, Edoardo Scopini, Gaia Grandin, Gabriele Cruciani, Antonio Giordano

## Abstract

**Objective:** Oxytocin (Oxt) assessment in plasma is challenging, and available data are contradictory. We aimed to assess circulating Oxt in mice by a validated nano-liquid chromatography/mass-spectrometry (nLC-MS/MS) protocol, combined with Oxt hypothalamic expression in different sex, life stages, and in diet-induced obesity.

**Methods:** We assessed plasma Oxt by nLC-MS/MS, *Oxt* hypothalamic expression by qPCR, and Oxt-immunoreactive neuron and fiber densities by immunohistochemistry and morphometric analyses in C57BL/6 mice at 21 and 60 days of life (p21 and p60, respectively). Mice in normo-fed condition and following 12 weeks of high-fat diet (HFD) were studied alongside food intake and hypothalamic expression of its regulators.

**Results:** Circulating Oxt does not vary based on sex at p21 and p60 but increases with aging. While hypothalamic *Oxt* mRNA expression followed the same trend across both sexes, Oxt neuron and fiber densities exhibited a similar trend only in females. Plasma vasopressin (Avp) followed Oxt trend in females but was opposite in males and was not mirrored by *Avp* mRNA hypothalamic expression. HFD-fed females were more resistant to weight gain compared to males and displayed higher Oxt plasma levels and hypothalamic expression. Sex dimorphism in food intake and hypothalamic expression of *Avp* and of key anorexigenic and orexigenic neuropeptides was detected.

**Conclusions:** Oxt plasma levels are higher in adulthood compared to weaning in mice of both sexes who displayed similar concentrations. Oxt plasma levels are mirrored by Oxt hypothalamic expression. In obesity, females display a lower increase in body weight but higher Oxt plasma levels than males.

**Significance:** This study delineates the pattern of Oxt plasma levels across various age, sex and in obesity in mice. We overcome some limitations of previous research, which were often confounded by numerous factors, by reliably measuring circulating Oxt and integrating this with parallel analyses of *Oxt* gene expression, and Oxt neuron and immunoreactive fiber densities. Critically, this is among the few studies to employ LC-MS/MS specifically in mice. This novel, integrated perspective is relevant given the compelling evidence for Oxt’s role as an anti-obesity agent and its potential as a therapeutic target. A precise understanding of Oxt regulation in different conditions, backed by reliable methodologies, is indispensable for the rational development of effective treatments for obesity and related endocrine diseases.

## 1. Introduction

Oxytocin (Oxt) is a neurohormone primarily produced by magnocellular and parvocellular neurons located in the hypothalamic paraventricular (PVN) and supraoptic (SON) nuclei. Oxt plays a pivotal role in the regulation of numerous behaviors, including sexual, reproductive, emotional, social, eating, and addictive behaviors. Most hypothalamic Oxt neurons project to the neurohypophysis where they release the neurohormone into the bloodstream upon various stimuli, most notably labor and lactation [1]. Nevertheless, peripheral Oxt exerts a wide variety of functions, among which the regulation of metabolic homeostasis [2]. The Oxt receptor is in fact widely distributed in metabolically relevant peripheral organs, including adipose tissues, liver, and pancreas [1]. Oxt administration not only exerts a central anorexigenic effect partly mediating leptin action [3], but also improves insulin sensitivity [4], induces lipolysis in white adipose tissue [5] and promotes thermogenesis in brown adipose tissue [6]. Consequently, Oxt has attracted significant interest for its potential action against obesity, leveraging both its central anorexigenic effect and its peripheral role in increasing energy expenditure [2].

However, Oxt plasma levels in physiological conditions across different life stages and sexes, as well as in conditions of obesity, remain controversial, primarily due to technical limitations related to its measurement [2]. The quantification of endogenous Oxt in body fluids has in fact represented a major concern in both human and animal studies. Oxt is composed of only 9 amino acids and is often bound to interfering plasma proteins, both aspects that make its detection highly challenging [7, 8]. To yield reliable Oxt concentration in mammalian blood, plasma samples need to be extracted to eliminate interfering proteins [7, 9]. Oxt concentration in unextracted plasma samples was shown to be 100-1000-fold higher than that in extracted plasma, levels that reflected the incorrect detection of substances other than Oxt [7]. The use of inappropriate and questionable detection assays strongly contributes to uncertainties among Oxt researchers and to the presence of inconsistent data in the literature. For example, while some studies report higher plasma Oxt levels among women compared to men, others show the opposite [10, 11]; similarly, it is still uncertain whether obesity is associated with higher or lower Oxt in circulation, as available studies often contradict each other [2] (Table 1). To increase the complexity of Oxt blood assessment there are numerous confounders known to affect its expression and circulating levels: stress, fear, social factors [12, 13], feeding/drinking [14, 15], circadian rhythm [16], estrous phase in females [17] can all contribute to the variability of Oxt levels. Lastly, a further possible confounder is represented by the close homology between Oxt and Vasopressin (Avp) molecules and systems [1].

**Table 1.** Circulating oxytocin levels in obesity across human and rodent studies.

| Species | Sex | Method Matrix | Obesity /<br>Normoweight | Notes | Reference |
| --- | --- | --- | --- | --- | --- |
| **Human**<br>**n=32** | 71,8%<br>Females | RIA and LC/MS<br>Plasma (extracted) | >1 | Reduced after bariatric surgery although<br>still higher than controls | Stock et al., Int J Obes 1989 [36] |
| **Human**<br>**n=176** | 64,7%<br>Females | ELISA<br>Serum (unextracted) | <1 | Lower among subjects with type 2 diabetes<br>vs those with normoglycemia | Qian et al., JCEM 2014 [26] |
| **Human**<br>**n=109** | 100%<br>Females | EIA<br>Plasma (extracted) | <1 | Lower with menopause | Maestrini et al., Ob Facts 2018 [37] |
| **Human**<br>**n=109** | 52,2%<br>Females | ELISA<br>Serum/plasma (extracted) | >1 | Increase across increasing BMI categories | Pataky et al., Int J Obes 2019 [25] |
| **Human**<br>**n=721** | 54,9%<br>Females | ELISA<br>Plasma (extracted) | >1 | Increase across increasing BMI categories | Weingarten et al., JCEM 2019 [38] |
| **Human**<br>**n=99** | 76,8%<br>Females | ELISA<br>Plasma (unextracted) | <1 | Lower levels only among metabolically<br>unhealthy subjects | Anvarova et al., Front Endocrinol 2025<br>[39] |
| **Mice** | Males<br>and<br>Females | EIA<br>Plasma (extracted) | =1 | Dietary protocol duration: 8 weeks for<br>males and 12 weeks for females | Maejima et al., Sci. Rep. 2017 [40] |
| **Mice** | Males | ELISA<br>Serum (extracted) | =1 | Dietary protocol duration: 10 weeks | Hayashi et al., Front. Endocrinol. 2020<br>[41] |
| **Mice** | Males | EIA<br>Serum (unextracted) | <1 | Dietary protocol duration: 4 months | Zhang et al., Neuron 2011 [14] |

The employment of specific and sensitive Oxt assays for a reliable quantification of the nonapeptide represents a central objective for Oxt research. Despite their growing commercial availability, quick performance and low cost, ELISA test reliability is critically dependent on the specificity of the antibody employed [8]. In contrast, Tandem Liquid Chromatography-Mass Spectrometry (LC-MS/MS) has been demonstrated to be the most reliable technique for successfully assessing Oxt quantities [2], although, to our knowledge, its application in mouse models has hitherto remained unexplored. In this study, we defined Oxt plasma levels across different life stages, sexes, and obesity states in mice, as assessed by a newly validated nano LC-MS/MS (nLC-MS/MS) protocol, thereby excluding most of the above listed confounders. To strengthen our results, we also evaluated whether the pattern of these plasma levels was mirrored by hypothalamic *Oxt* mRNA expression, by the densities of Oxt neurons and of Oxt-immunoreactive (Oxt-IR) fibers. Lastly, we examined Oxt levels in conjunction with feeding behavior and the expression of key hypothalamic energy balance regulators to identify possible sex-dimorphic responses to obesity.

## 2. Materials and Methods

### 2.1 Animal housing and dietary protocols

Female and male C57BL/6 mice were purchased from Charles Rivers (Calco LC, Italy), housed in plastic cages and maintained on a 12:12h light-dark cycle at 22-23 °C with free access to standard diet (Mucedola srl., Settimo Milanese, MI, Italy #4RF21) and water. All experimental groups were composed of at least n=5 animals for plasma and gene expression assessments and of n=3 animals for morphological analyses, consistently with morphometric studies. All females studied were virgin. Animals were sacrificed at post-natal day (p) 21, time of weaning, at p60 (early adulthood), and at 4 months of age (p120) in the obesity experiment as described below. All animal procedures were performed in accordance with the national ethical guidelines and with the approval of the local animal care and use committee *Organismo Preposto al Benessere Animale* and of the *Istituto Superiore di Sanità*, authorization number 15/2023-PR, in accordance with the law art. 31 of D.lgs. 26/2014.

In the diet-induced obesity experiment, four-week-old animals were housed in groups of two mice per cage and allowed to acclimatize for 1 week, when they were fed with the above standard diet (week 0). Animals were not housed singularly (two per cage) to avoid any interference of stress and loneliness on the Oxt system [13]. Animals were then divided into 2 experimental groups: one group fed a chow diet (CD) (10% fat, 70% carbohydrates, 20% proteins; Research Diets, New Brunswick, NJ; #D12450K), while the second group fed a high-fat diet (HFD; 45% fat, 35% carbohydrates, 20% proteins; Research Diets; New Brunswick, NJ #D12451). Each cage was provided with the same amount of diet (70g per week). These feeding conditions were maintained for 12 weeks. Animals and residual food were weighed once a week to monitor body weight trends and food intake, respectively. Food intake is expressed as kilocalories (kcal), reported either as absolute values or normalized to body weight. Food intake percentage change data were adjusted for body weight. Animals participating in this experiment were sacrificed after conclusion of the dietary protocol at p120 when samples were collected for the downstream analysis.

### 2.2 Sample processing

All actions taken to minimize confounding factors are listed in Table 2. In brief, to avoid the impact of circadian Oxt fluctuation, all samples were collected between 2:00-4:00 PM, when Oxt levels are more stable [16]. To ensure the lack of restraint-evoked stress response which could affect Oxt circulating levels or hypothalamic expression factors [12], one month before their sacrifice, mice were handled three times per week. To avoid the sacrifice during the estrus phase, the week before the start of the experiment female mice were observed once daily to assess the estrous cycle. The stage of the estrous cycle was determined by a non-invasive visual assessment method, chosen as an alternative to the vaginal smear technique. This method is as accurate and reliable as the vaginal smear, and it is non-stressful to animals [18]. To minimize variability, all assessments were performed by a single-trained operator. Four hours before the sacrifice, food was removed to reproduce consistent conditions across animals and minimize the impact of food intake. Animals underwent isoflurane anesthesia before the sacrifice, a technique chosen for being minimally stressful and invasive for the animal.

**Table 2.** Measures Implemented to Control Confounding Factors in Plasma Oxytocin Measurement.

| CONFOUNDING FACTOR | HOW IT AFFECTS MEASUREMENT | ACTIONS TAKEN |
| --- | --- | --- |
| STANDARDIZED CONDITIONS BEFORE SAMPLE COLLECTION |  |  |
| DIURNAL FLUCTUATIONS [16] | Natural variability masks differences | Collect at consistent times; repeated measures |
| STRESS [12] | Stress increases endogenous release | Minimize discomfort; regular animal handling |
| DIET, HYDRATION [14, 15] | Influences hormone levels indirectly | Standardized fasting and food intake |
| ESTROUS CYCLE [17] | Strong biological variability | Estrous cycle phase assessed to standardize collection timing |
| SOCIAL ISOLATION [13] | Reduces oxytocin levels | Mice were housed at a minimum of two per cage |
| SAMPLE COLLECTION AND PROCESSING |  |  |
| RAPID DEGRADATION BY PEPTIDASES [42] | Leads to artificially low concentrations | Keep samples on ice; rapid processing |
| CHOICE OF ANTICOAGULANT | Affects peptide stability and binding | Use validated anticoagulant - EDTA |
| CROSS-REACTIVITY IN IMMUNOASSAYS [8] | Overestimation due to similar peptides | Use of the highly specific LC–MS/MS assay |
| MATRIX EFFECTS [8] | Interference with assay sensitivity | Perform extraction; validated sample prep |
| EXTRACTION VARIABILITY [43] | Affects final concentration | Standardize protocols; assess apparent recovery (internal standardization) |
| STORAGE CONDITIONS | Degradation over time | Store at –80°C; avoid freeze–thaw |

Blood for plasma Oxt was collected by intracardiac puncture under anesthesia, followed by decapitation of the animal for hypothalamus retrieval destined to gene expression analyses. In brief, blood was placed in 1.5 mL-tubes containing 100 μl of EDTA (0.5M, pH8) and immediately stored in ice. Samples were quickly processed for plasma collection which occurred by centrifugation at 3000 rpm for 10 min using a refrigerated (4°C) centrifuge. The skull of the animal was carefully opened, and the whole hypothalamus (approximately 5 mm thick) was isolated using a brain matrix and surgical blades. The accuracy of the collection procedure was verified in a separate set of animals by Nissl staining to confirm the correct isolation of the PVN, SON, and arcuate (ARC) nucleus. The resulting hypothalamic samples were immediately frozen in liquid nitrogen and stored at –80°C until RNA extraction.

For the morphological analyses, another set of mice was perfused transcardially with 4% paraformaldehyde in 0.1M phosphate buffer saline (PBS), pH 7.4. Brains were carefully removed from the skull, post-fixed with the same fixative solution for 24h at 4°C and washed in PBS. Hypothalamus isolation was performed as described above. Free-floating coronal sections (40 μm thick) were then obtained with a Leica VT1200S vibratome (Leica Microsystems, Vienna, Austria) and kept in PBS, pH 7.4, at 4°C until use.

### 2.3 Oxytocin and vasopressin quantification in plasma by nLC-MS/MS

Oxt was assessed according to human sample (urine and serum) analysis procedure of Franke et al. [19] adapted to and validated for murine plasma analysis as described in Grifnée et al. [20]. Samples were prepared for analysis adding 5 μL of the internal standard oxytocin-(leucine-5,5,5-d3, glycine-2,2-d2) trifluoroacetate salt (Merck, Burlingtone, MA; #900169-1MG; at 10 μg/L) to 100μL of plasma. To create a calibration curve, oxytocin acetate salt hydrate ≥97% purity for HPLC (Merck; #O6379) was used. Samples were added with 120 μL of H_3_PO_4_ and shaken for 45 min at room temperature (RT). After the addition of 600 μL of acetonitrile (ACN), samples were kept at -20°C for 30 min and then centrifuged at 14000 rpm for 10 min at 4°C. Supernatant was transferred in a tube and evaporated to approximately 300 μL at 45°C and under a gentle stream of nitrogen. After adding 3 mL of water, samples were purified by the solid-phase extraction (SPE) columns Strata-X 33 μm Polymeric Reversed Phase 60 mg/3 mL (Phenomenex, Torrance, CA; 8B-S100-UBL). SPE columns were conditioned with 3 mL of methanol and 3 mL of water. Samples were loaded into the columns and were first washed with 3 mL of water and then with 3 mL of ACN 5%. Analytes were eluted with 3 mL of ACN 70%, and solutions were evaporated till dryness in evaporating centrifuge. Residues were resuspended in 200 μL of ACN 10% containing 0.1% of formic acid (FA) and transferred to a new 2 mL tube. After the addition of 700 μL of ACN, samples were kept at -20°C overnight and then centrifuged at 14000 rpm for 10 min at 4°C. The supernatant (850 μL) was evaporated under a gentle nitrogen stream at 45°C and residues were suspended in 100 of ACN 10% containing 0.1% of FA. As quality control, a pooled plasma sample was treated as previously described adding 5 µL of oxytocin at 10 µg/L before the extraction. The nanoLC coupled to a Q-Exactive PLUS system (Thermo Fisher Scientific, Waltham, MA) was used for the analyses. Acclaim PepMap RSLC 75 μm x 25 cm column (Thermo Fisher Scientific, Waltham, MA) was used for chromatographic separation employing H_2_O and ACN, both containing 0.1% of FA as mobile phase A and B respectively. Gradient initiated with 2% of mobile phase B maintaining this condition for 2 min and then the mobile phase B increased to 30% in 63 min. The final step increased the percentage of mobile phase B to 72% in 12 min and this condition was maintained for 6 min. The flow rate was 0.3 μL/min, column temperature and samples were set at 40°C and 16°C respectively. 5 μL of samples were injected using trap-elute based approach. Before the injection on the analytical column, PepMap NEO C18 5µm, 300 µm x 5 mm trap column (Thermo Scientific 174500) was used to trap the analytes using water with 0.1% of formic acid as mobile phase at flow of 30 µL/min for 3 min. Nano electrospray ionization (nESI) operating in positive polarity and target-SIM/dd-MS2 experiments were used for the analyte’s ionization and detection. Resolution, AGC (Automatic Gain Control) target, maximum injection time and isolation window was set at 140000 (FWHM@200 m/z), 1e6, 400 ms and 2.0 m/z respectively for full scan experiment. For dd-MS2 experiment resolution, AGC targeted, maximum injection time, isolation window, Normalized Collision Energy (NCE), minimum AGC target, apex trigger and dynamic exclusion were set 35000 (FWHM@200 m/z), 5e5, 150 ms, 1 Da, 10-20-35 eV, 8e2, 1-3 s and 2 s respectively. For the detection of the analyte and the internal standard, the *m/z* values 504.2255 and 506.7412 were monitored, corresponding to doubly charged oxytocin and doubly charged oxytocin-d5.

### 2.4 Vasopressin quantification in plasma by nLC-MS/MS

After addition of 5 μL of oxytocin-(leucine-5,5,5-d3, glycine-2,2-d2), 100 µL of plasma sample were incubated under agitation with 120 µL of 4% H_3_PO_4_ solution in 2 mL low protein binding tubes at RT and in the dark. After 45 min of incubation, 600 µL of cold ACN was added and the samples were vortexed and placed at -20°C for 30 min for protein precipitation. Then, samples were centrifuged for 10 min at 14000 rpm and 4°C. The supernatant was transferred into 15 mL tubes and was evaporated under nitrogen at maximum temperature of 45°C to remove organic solvent. 3 mL of 4% H_3_PO_4_ solution was added, and the analytes were extracted with Waters Oasis WCX (150 mg/6 mL) columns (Waters Corporation, Milford, MA). The samples were loaded on the columns after the conditioning with 6 mL of CH_3_OH and 6 mL of H_2_O and the columns were washed with 5 mL of 4% NH_4_OH solution and 5 mL of 10% CH_3_OH solution. The analytes were eluted with 3 mL of 90% of ACN solution with 1% of trifluoroacetic acid and the elute was recovered in 5 mL Eppendorf tubes. Then, samples were evaporated to dryness in an evaporating centrifuge at 45°C and were regained with 100 µL of 5% CH_3_OH solution containing 0.1% of HCOOH. The samples were filtered with LCR filters (Merck; #SLLGR04NL) and analyzed in Thermo nanoLC-Q-Exactive PLUS instrument. For the analysis, the chromatographic and spectrometric conditions were the same as those employed for oxytocin detection. For the detection of Avp, the *m/z* value 542.7262 was monitored, corresponding to doubly charged vasopressin.

### 2.5 nLC-MS/MS method validation

Method validation was performed by spiking a pooled plasma sample at 2.5 ng/L, 10 ng/L, 50 ng/L, 100 ng/L, 500 ng/L and 1000 ng/L for Oxt and 25 ng/L, 50 ng/L, 100 ng/L, 500 ng/L and 1000 ng/L for Avp. Each level was analyzed in four replicates over three different days preparing each sample as reported above.

#### Selectivity

Samples fortified with both oxytocin and vasopressin were prepared according to the method extraction and injected into nUHPLC-HRMS to confirm the chromatographic separation between other compounds naturally presented in the matrix. Moreover, the obtained MS/MS spectra were compared to confirm the characteristic fragmentation patterns.

#### Solvent linearity

Solvent linearity was verified by injecting three times each standard at 2 ng/L, 5 ng/L, 10 ng/L, 25 ng/L, 50 ng/L, 100 ng/L, 250 ng/L, 500 ng/L and 1000 ng/L for oxytocin and 20 ng/L, 50 ng/L, 100 ng/L, 250 ng/L, 500 ng/L and 1000 ng/L for vasopressin all containing oxytocin-d5 at 500 ng/l and reporting in y-axis the area ratio oxytocin/oxytocin-d5 or vasopressin/oxytocin and in x-axis the oxytocin or vasopressin concentrations. Linearity was considered acceptable in the range where difference of back calculated concentration was less than 10%.

#### Matrix linearity

Matrix linearity was verified by injecting the same concentration points injected for the solvent linearity assessment using undiluted plasma for higher levels (25 ng/L, 50 ng/L, 100 ng/L, 250 ng/L, 500 ng/L and 1000 ng/L for oxytocin and 100 ng/L, 250 ng/L, 500 ng/L and 1000 ng/L for vasopressin) and diluted plasma (1:5) for lower levels. Linearity was considered acceptable in the range where the deviation of back-calculated concentrations was less than 15%.

#### Trueness and precision

Trueness and precision were evaluated using undiluted pooled plasma for the higher concentration ranges (25 ng/L, 50 ng/L, 100 ng/L, 500 ng/L and 1000 ng/L for oxytocin and 100 ng/L, 250 ng/L, 500 ng/L and 1000 ng/L for vasopressin), whereas a 1:5 pre-extraction dilution was applied for the lowest levels (2.5 and 10 ng/L for oxytocin and 25 and 50 ng/L for vasopressin). Trueness, intraday precision (repeatability) and inter-day precision (within-laboratory reproducibility) for each validation level, pooled apparent recovery (%), pooled repeatability (CVr, pooled (%)) and pooled within-laboratory reproducibility (CVR, pooled (%)) are reported in Table S1.

#### Limit of quantification and uncertainty

Limits of quantification (LOQ) were set at lowest validation level for both oxytocin (2.5 ng/L) and vasopressin (25 ng/L). Relative combined uncertainty (ucomb%) was calculated considering the contribution of CVR, pooled (%), coefficient of variation (CV%) of recovery, the contribution of calibration considering rectangular distribution and 1 replicate for each real sample. The obtained values were 15.6% and 15.5% for oxytocin and vasopressin respectively. In Fig. 1C e 1D, blank, standard at 250 ng/L and real samples at low concentration along their obtained MS/MS spectra for both Oxt and n are shown.

**Figure 1.**
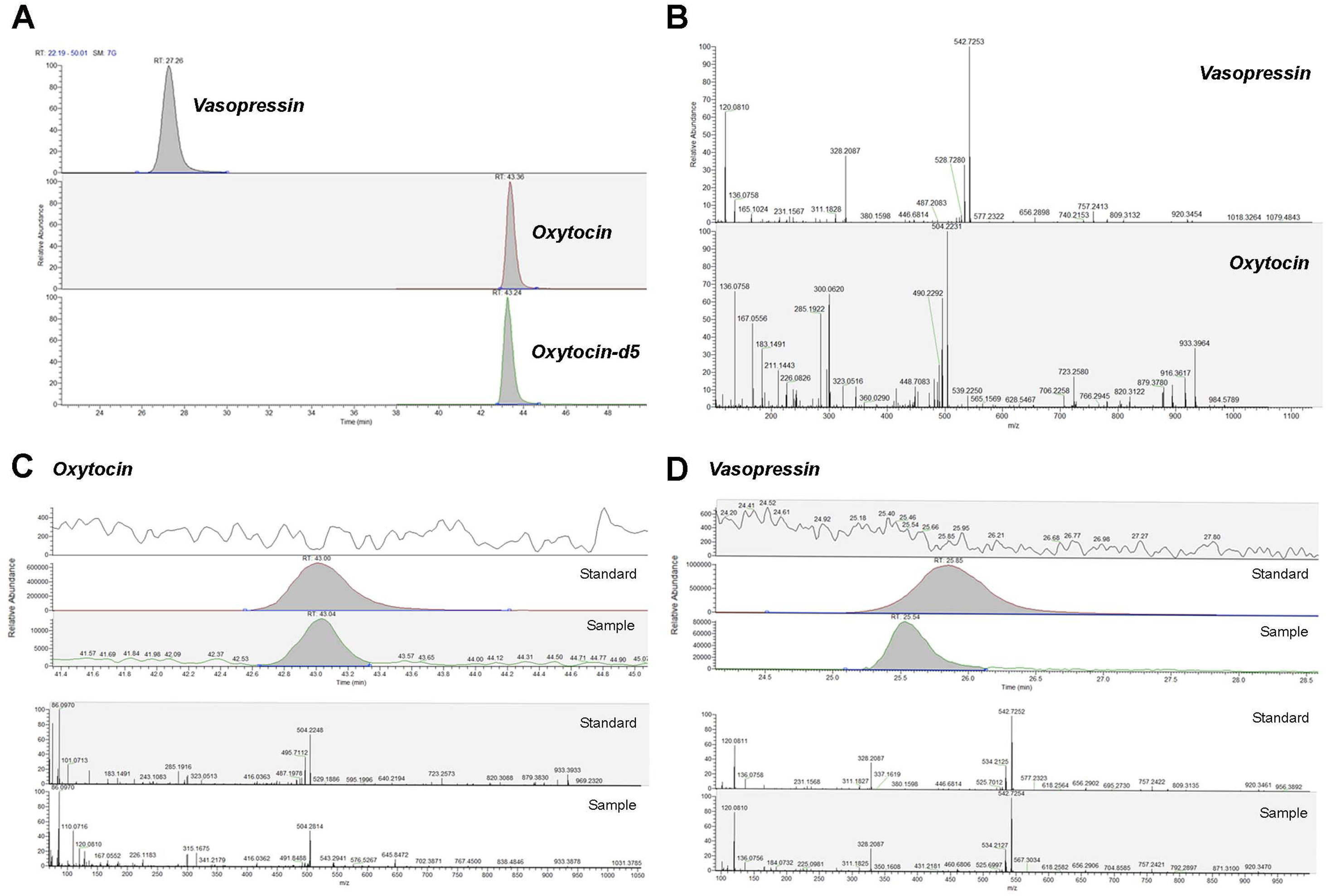
Analytical validation of the nLC-MS/MS procedure for Oxytocin and Vasopressin quantification. Representative nLC-MS/MS chromatogram, showing three distinct peaks: the target analyte oxytocin, the structurally similar nonapeptide vasopressin, and the isotopically labeled internal standard (Oxytocin-d5) (**A**). High-resolution MS/MS fingerprints for Vasopressin (top) and Oxytocin (bottom), matching the fragmentation patterns of authentic standards to ensure unequivocal analyte identification (**B**). Comparison of chromatographic peaks and MS/MS spectra between standard solutions (250ng/L) and representative plasma samples at the LOQ level (2.5 ng/L for Oxytocin; **C**; 25 ng/L for Vasopressin; **D**)

### 2.6 Hypothalamic gene expression by qPCR

RNA extraction was performed using phenol-chloroform-guanidinium thiocyanate method. Briefly, each sample was mixed with 500 μl of Trizol (Invitrogen, Waltham, MA) and a sterile and RNAse free zirconium grinding bead was placed in each tube to perform tissue homogenization by TissueLyser (Qiagen, Hilden, Germany) for 5 min at 50 Hertz. Samples were added with chloroform and then centrifuged for 15 min at 12000 rpm at 4°C. An equal volume of 70% ethanol was added to the isolated supernatant. RNA extraction was performed using the Total RNA Purification Kit (Norgen Biotek Corp., Thorold, ON, Canada; #17250) combined with the RNase-Free DNase I Kit (Norgen Biotek Corp., #25720) to remove genomic DNA. The quality and quantity of total RNA was evaluated using a Nanodrop Spectrophotometer (Fisher Scientific, Hampton, NH). One μg RNA was reverse-transcribed with the High-Capacity cDNA RT Kit with RNase Inhibitor (Applied BioSystems, Waltham, MA; #4374966) in a total reaction volume of 20 μl. The cDNA library was then used for quantitative real-time PCR. For SYBR® Green gene expression assays: pairs of primers were designed by Primer3 (web version 4.1.0) and the NCBI Primer Blast software, and then customized by ThermoFisher (Waltham, MA; # 10336022) (Table S1). Universal SYBR Green Master Mix (Applied Biosystem, Foster City, CA; #4309155) was employed for the cDNA amplification. All reactions were carried out by the Step One Plus Real-Time PCR system (Applied BioSystems) using 50 ng RNA on a final reaction volume of 10 μl. The thermal cycling conditions for SYBR® Green assays followed the standard cycling mode for designed primers with a Tm ≥60: 1 cycle at 50 °C for 2 min, 1 cycle at 95 °C for 2 min and 40 cycles at 95 °C for 3 sec followed by 60 °C for 30 sec. A dissociation step (melt curve stage) of 95°C for 15 sec, 60 °C for 1 min and 95 °c for 15 sec, was included at the end of the cycling stage. All samples were run in duplicate. Samples not containing the template were included as negative controls in all experiments. β-actin was used as endogenous controls to normalize gene expression. Relative mRNA expression was determined by the ΔΔCt method (2^−ΔΔCt^).

### 2.7 Immunohistochemistry, confocal microscopy, and morphometric analyses

Immunohistochemical detection of Oxt was performed according to standard procedures. Briefly, free-floating sections reacted with 1% NaOH and 1% H_2_O_2_ (20 min), 0.3% glycine (10 min), and 0.03% sodiumdodecylsulfate (10 min) in PBS. After rinsing in PBS, they were blocked with 3% normal goat serum (Vector Laboratories, Burlingame, CA; #S-37 1000-20) in 0.2% TritonX-100 in PBS for 60 min and incubated with the primary antibodies (Table S2) in PBS overnight at 4^°^C. The next day, after a thorough rinse in PBS, sections were incubated in 1:200 biotinylated secondary antibody (Table S2) in PBS for 30 min at RT, rinsed in PBS, and incubated in avidin-biotin peroxidase complex (ABC Elite, Vector Laboratories; #PK-6100) for 1h at RT. After washing several times in PBS, sections were finally incubated for 1 min in 3,3’-diaminobenzidine solution (Vector Laboratories; #SK-4100) at RT. After the staining, sections were mounted on slides, air-dried, dehydrated in ethanol, cleared with xylene, and mounted with Eukitt (Bio-Optica S.p.A, Milan, Italy). Reactions were analyzed using a Nikon Eclipse E600 microscope (Nikon; Sesto Fiorentino, Florence, Italy) with a camera mounted to obtain the images. Pictures were taken at a magnification of 20x and 40x, and brightness and contrast of the final images were adjusted using Photoshop 6 software (Adobe Systems, Mountain View, CA). For immunofluorescence and confocal microscopy experiments, free-floating sections were processed according to the above-described protocol for immunohistochemistry up to incubation with the primary antibodies. The next day, after 3 washes in PBS and additional incubation with 4% normal donkey serum in PBS (Jackson ImmunoResearch; Bar Harbor, MA; # 017-000-001) for 30 min at RT, sections were incubated in fluorophore-linked secondary antibodies (Table S2) solution in PBS for 1h at RT. Nuclear staining was performed using the fluorescent dye TO-PRO-3 Iodide (642 nm of excitation wavelength) (Invitrogen by Thermo Fisher, MA; #T3602), in PBS for 15 min at RT. Sections were subsequently washed twice with PBS, mounted on standard glass slides, air-dried, and covered using Vectashield mounting medium (Vector Laboratories; # H-1000-10). Samples were analyzed by a motorized Leica DM6000 microscope at 40X and 60X magnifications. Fluorescence was detected with a Leica TCS-SL spectral confocal microscope. Fluorophores were excited with 488, 543, and 649 nm lasers and imaged separately. Images (1024 × 1024 pixels) were obtained sequentially from two channels using a confocal pinhole of 1.1200. The brightness and contrast of the final images were adjusted using Photoshop6 (AdobeSystems).

To minimize procedural variability, morphometry was performed on sections from different experimental groups that were stained in parallel. The quantification of Oxt neurons in PVN and SON was performed on immunofluorescent stained sections by manually counting the number of neurons belonging to each nucleus. Specifically, we summed the number of Oxt neurons belonging to at least three non-overlapping coronal sections per mouse representing the rostral, medial, and caudal hypothalamic PVN and SON at 40x magnification. Data are expressed as the average number of Oxt neurons per slide. Oxt-IR fibers in the PVN and SON was performed following the protocol described by DiBenedictis et al [21] on immunofluorescent-stained slices. Brain sections containing the PVN and SON were imaged using a 20x objective. Then, Oxt-IR fibers were measured by manually thresholding the gray-scale images using the ImageJ software (NIH; imagej.nih.gov/ij). To ensure that the Oxt-IR fiber density measurement specifically reflected fiber density and caliber, Oxt cell bodies present within the sampling area were removed from the calculation. The pixels attributed to these cell bodies were excised manually using the polygon draw tool in ImageJ. As described in DiBenedictis et al. this analysis provides an estimate of the fractional area occupied by Oxt-IR elements, which is considered to reflect the staining intensity, fiber density, and fiber caliber/thickness. Oxt-IR fiber density data are reported as the number of pixels within each section where Oxt immunoreactivity was detected, with a greater number of pixels indicating a larger proportion of the area occupied by Oxt elements.

### 2.8 Statistical analysis

All data are reported as mean ± standard error of the mean (SEM) in graphs. Comparisons between groups were performed with Student’s t-test. Pearson’s correlation coefficient (r) was calculated to assess the linear relationship between variables. A p < 0.05 was considered statistically significant. All statistical analyses were performed with GraphPad Prism 6 (GraphPad Software, San Diego, CA) software and RStudio (Integrated Development Environment for R. Posit Software, PBC, Boston, MA).

## 3. Results

### 3.1 Analytical performance and validation of the nLC-MS/MS procedure for oxytocin and vasopressin quantification

To ensure the accuracy and reliability of neuropeptide quantification, we performed method validation in mouse plasma. The method demonstrated high selectivity, with a clear chromatographic separation of Oxt and Avp from endogenous matrix interferences (Fig. 1A). The identity of the target analytes was confirmed by matching fragments of authentic standards and analytes in biological samples (Fig. 1B). Detailed structural characterization of Oxt was further supported by the identification of a diagnostic fragment at m/z 300.0614 ([C_14_H_11_O_5_N_3_] ^+^), originating from the rearrangement of the cyclic moiety (Fig. S1A-B). The assay exhibited high linearity in both solvent (R^2^ > 0.99) and plasma (R^2^ > 0.99) samples over the investigated concentration ranges (2–1000 ng/L and 25–1000 ng/L for Oxt and Avp, respectively). The high reliability of the quantitative data was further supported by the trueness and precision parameters shown in Supplementary Table S3A-B. The method showed high accuracy across all validation levels, with pooled apparent recoveries of 100% and 103% for Oxt and Avp, respectively. Furthermore, the assay proved to be highly precise; notably, at the most challenging concentrations, the intraday precision remained high, with CV% values as low as 4.1% for Oxt (at 2.5 ng/L) and 8% for Avp (at 25 ng/L). The method’s sensitivity was confirmed by the detection of clear, distinct peaks at the LOQ levels (2.5 ng/L for Oxt, Fig. 1C: 25 ng/L for Avp, Fig. 1D). The MS/MS spectra obtained from plasma samples at these concentrations showed a high match with the fragmentation patterns of authentic standards, demonstrating sensitivity levels for this analytical method comparable to previously published procedures [20].

### 3.2 Oxytocin plasma levels and hypothalamic expression are higher in male and female adult compared to weaning mice

Plasma Oxt is released by hypothalamic magnocellular oxytocinergic neurons of the PVN and SON, which, in contrast to parvocellular neurons, project to the neurohypophysis and directly contribute to circulating Oxt levels (Fig. 2A and B) [22]. Oxt plasma levels did not vary based on sex but were higher at p60 compared to p21 in both females and male mice (Fig. 2C). We hence corroborated our findings assessing *Oxt* mRNA hypothalamic expression, which in fact followed the same trend of plasma Oxt levels, with significantly higher *Oxt* in male and female older animals (Fig. 2D). Synaptotagmin 4 (Syt4) is a negative regulator of synaptic exocytosis, mostly expressed by Oxt neurons [14]. To assess whether differences in Oxt plasma concentrations could be associated with a variation in the levels of its regulator of exocytosis, we also assessed *Syt4* mRNA hypothalamic expression. *Syt4* expression paralleled that of hypothalamic and plasma Oxt levels (Fig. 2E). In this context, unlike conditions of acute stimulation of Oxt secretion, an inverse relationship between Syt4 and Oxt is not necessarily expected, as Syt4 acts as a regulator of Oxt release; therefore, higher levels of Oxt may require a corresponding increase in the regulatory machinery involved in its exocytosis. Notably, Syt4 function appears to depend primarily on its vesicular localization and involvement in exocytotic regulation rather than on transcript abundance per se [14].

**Figure 2.**
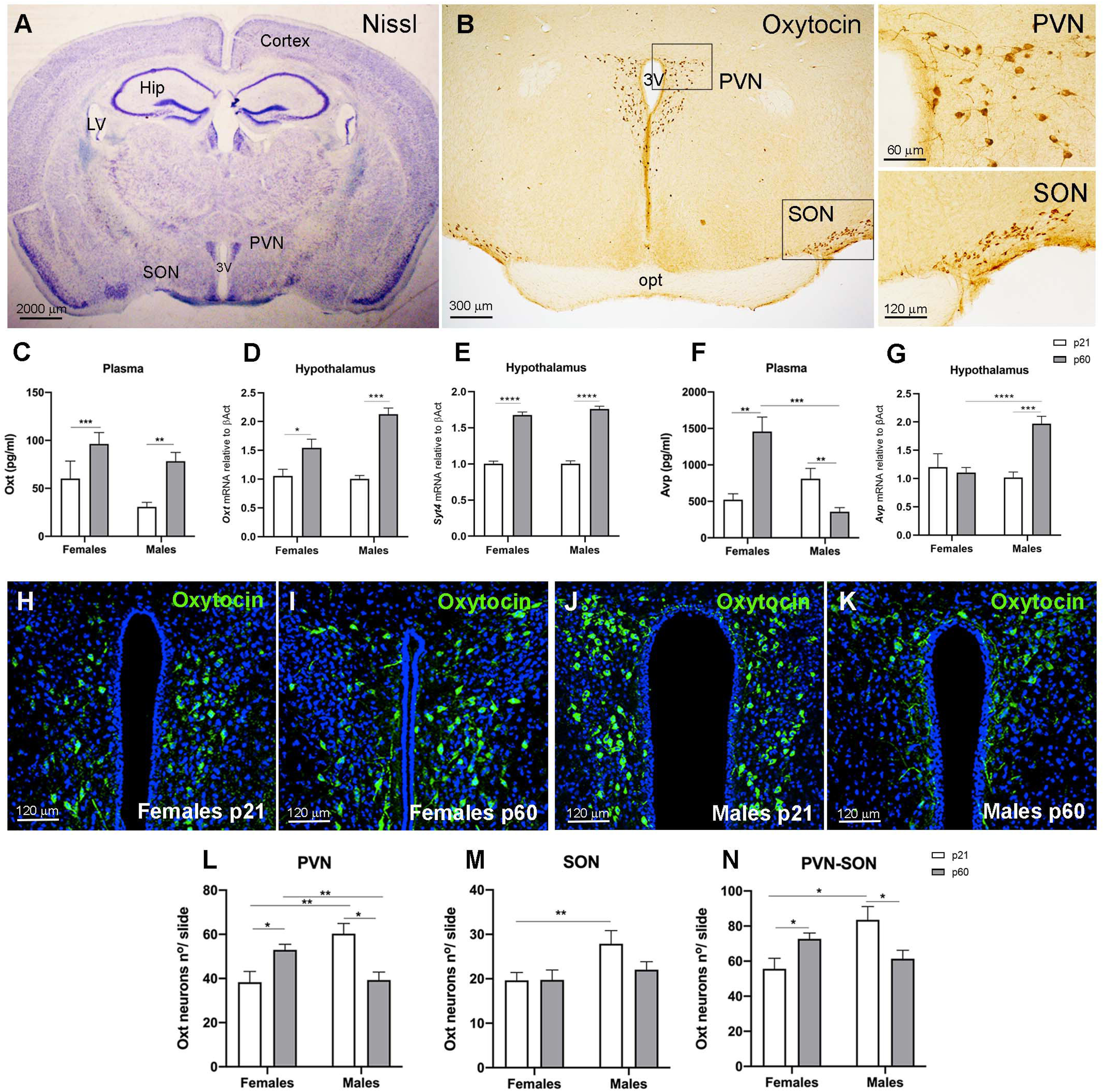
Oxytocin plasma levels, hypothalamic expression, and neurons number across different ages and sexes in mice. Light microscopy of Nissl-stained brain coronal section (**A**) and Peroxidase immunohistochemistry for Oxytocin (Oxt) (**B**) of the region corresponding to bregma −0.94 mm. Enlargements in B are representative of the squared area, depicting Oxt-immunoreactive neurons and fibers in the PVN and SON. The following outcomes were compared in female and male mice at 21 and 60 days of life moment of weaning when mice are still breastfed, and of early adulthood (p21 and p60, respectively): Oxt plasma levels (**C**); hypothalamic *Oxt* (**D**) and Synaptotagmin-4 (*Syt4)* (**E**) mRNA expression; Vasopressin (Avp) plasma levels (**F**); hypothalamic *Avp* mRNA expression (**G**). Representative images of Oxt-immunoreactive neurons in the PVN of females at p21 (**H**) and p60 (**I**) and of males at p21 (**J**) and p60 (**K**), by immunofluorescence and confocal microscopy. All nuclei were stained in blue by TO-PRO3. Total number of Oxt-immunoreactive neurons per slice in the PVN (**L**), in the SON (**M**) and in the combined Oxt neurons population (PVN and SON) (**N**) by sex and age. 3V= third ventricle; PVN: hypothalamic paraventricular nucleus; SON: hypothalamic supraoptic nucleus; LV=lateral ventricle; Hip: hippocampus; opt= optic tract. Comparisons between groups were performed with Student’s t-test. Data are expressed as mean ± SEM, *p<0.05; **p<0.01; ***p<0.001. N=minimum 5 mice per group for plasma and gene expression analysis; N=3 mice per group for morphometric analysis.

Avp is a neuropeptide structurally closely related to Oxt, also produced by PVN and SON, and released through the neurohypophysis [1]. To verify the accuracy of our methodology and assess whether these nonapeptides followed the same trend, Avp was also measured in plasma. nLC-MS/MS revealed distinct chromatographic separation of Oxt and Avp: despite their structural similarity and low molecular weight, the two hormones exhibited significantly different retention times, resulting in well-resolved peaks on the chromatogram (Fig. S1A). Avp plasma levels were higher at p60 compared to p21 in females, while the opposite trend was found in males who also displayed significantly lower Avp levels in adulthood compared to their female counterparts (Fig. 2F). Differently from *Oxt*, *Avp* mRNA hypothalamic expression did not follow the same trend as its plasma levels: it in fact remained stable during aging in females, while it increased in males who displayed significantly higher levels at p60 (Fig. 2G).

These data indicate that Oxt, but not Avp, plasma levels are paralleled by a similar hypothalamic mRNA expression.

### 3.3 Oxytocin plasma levels are associated with hypothalamic oxytocin neuron and fiber densities in females, but not in males

To assess whether the differences in Oxt plasma levels between p21 and p60 old mice were associated with a variation in the density of hypothalamic Oxt neurons, a morphometric study of PVN and SON was also performed. To identify the best antibody able to detect Oxt neuron bodies to perform morphometric analysis, two primary antibodies recognizing Oxt were tested (Table S2). The antibody from ImmunoStar was selected to study Oxt neuron density due to its ability to better detect Oxt neuron bodies (Fig. S1C and D). In the PVN of female mice, Oxt neuron density was higher at p60 compared to p21, similarly to Oxt plasma levels and hypothalamic expression (Fig. 2H-I, and L); however, male mice displayed a reduced Oxt neuron density with aging in the same area (Fig. 2J-K, and L). Overall, in the PVN, male mice displayed a higher Oxt neuron density compared to females at p21, while the last had the highest density of Oxt neurons at p60 (Fig. 2L). In SON, Oxt neuron density was higher in males at p21, but no other differences were detected in this area (Fig. 2M). Collectively, hypothalamic Oxt neuron density – assessed including both PVN and SON in the analysis – was higher at p60 compared to p21 in females but followed the opposite trend in males (Fig. 2N). This sexual divergence detected in adulthood may be related to the impact of sex-specific hormonal changes occurring during puberty. Part of PVN Oxt neurons extend their axons ventroposteriorly (Fig. 3A and B) to form the tuberohypophysial and hypothalamoneurohypophysial systems, last of which reaches the neurohypophysis where Oxt is released in the systemic circulation (Fig. 3A) [22]. Both systems cross the median eminence where Oxt axons form swellings of neurosecretes known as *Herring bodies* (Fig. 3C), also found in the posterior pituitary. To assess whether differences in Oxt plasma levels could be associated with variations in the density of axons of PVN Oxt neurons projecting ventroposteriorly to the median eminence and posterior pituitary, Oxt-IR fiber density was evaluated. The antibody from Synaptic System was selected for the analysis of Oxt-IR fibers due to its relative superior visualization of Oxt-IR fibers (Fig. S1C and D). Overall, Oxt-IR fiber density followed the same trend of Oxt neuron density with higher values in females at p60 compared to p21 and an opposite trend in males (Fig. 3D). In particular, PVN Oxt-IR fiber density in females was higher at p60, while it was lower at p21 compared to males (Fig. 3D). As SON Oxt neuron mostly project to the neurohypophysis where secrete the nonapeptide, the density of Oxt-IR fibers was also evaluated in this region. Oxt-IR fiber density in this area mirrored that of the PVN (Fig. 3E). Taken together, these data suggest that the differences in Oxt neuron and fiber densities are associated with similar differences in Oxt plasma levels only in female mice and were opposite in males. Based on this finding, the underlying morphological architecture supporting Oxt release follows a sexually dimorphic and opposite age-related trajectory.

**Figure 3.**
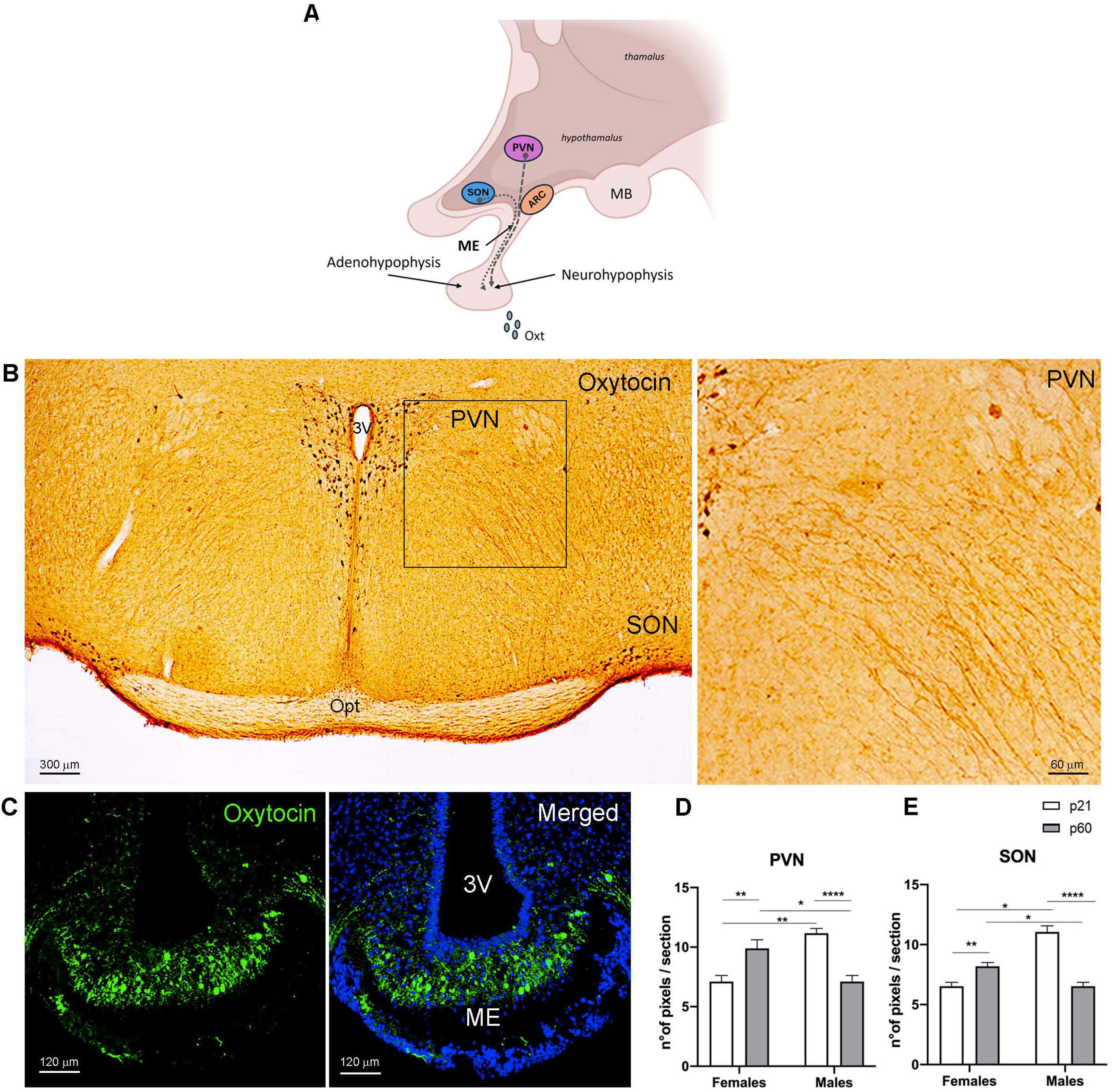
Oxytocin-immunoreactive fiber density across different ages and sexes in mice. Schematic representation (created via BioRender) of the hypothalamoneurohypophysial system. The diagram shows the course of oxytocinergic fibers originating from the PVN and the SON and projecting towards the neurohypophysis for systemic Oxt release (**A**). Peroxidase immunohistochemistry and light microscopy for Oxt in the PVN and the SON (**B**). The enlargement is representative of the squared area depicting an abundant cascade of Oxt-immunoreactive fibers descending ventrally from the PVN (**B**). Immunofluorescence and confocal microscopy for Oxt showing neurosecretory swellings, known as Herring bodies, within the median eminence, representing sites of Oxt storage and release (**C**). All nuclei were stained in blue by TO-PRO3. Morphometric analysis of Oxt-immunoreactive fibers in the PVN (**D**) and in the SON (**E**). 3V= third ventricle; PVN: hypothalamic paraventricular nucleus; SON: hypothalamic supraoptic nucleus; opt= optic tract; ARC= arcuate nucleus; MB= mammillary bodies; ME= median eminence; p= postnatal day. Comparisons between groups were performed with Student’s T-test. Data are expressed as mean ± SEM, *p<0.05; **p<0.01; ***p<0.001. N=3 mice per group for morphometric analysis.

### 3.4 Oxytocin plasma levels are higher in female compared to male mice with obesity

We subjected one month old male and female mice to a 12-week HFD or CD protocol to evaluate whether Oxt plasma and hypothalamic mRNA levels are altered following diet-induced obesity. Female mice fed a HFD were relatively resistant to weight gain, as their body weight curve overlapped with that of CD at almost all timepoints (Fig. 4A and C). On the other hand, male mice were more prone to obesity as they gained a significantly higher amount of weight when exposed to HFD compared to CD (Fig. 4B and C). Notably, while females rapidly compensated for the initial increase in caloric intake by reducing their overall energy intake, this adaptive response was less evident in males, which maintained higher caloric intake at week 2 and showed no consistent differences thereafter (Fig. 4D). This sex-dependent pattern of energy intake may contribute to the differential body weight gain observed between males and females. Consistently, food intake normalized to body weight was increased during the first week of HFD exposure in both sexes, likely reflecting an initial hypercaloric response in the absence of significant differences in body weight (Fig. 4E). On the other hand, in subsequent weeks, food intake normalized to body weight progressively declined in both males and females (Fig. 4E). While this reduction is consistent with the compensatory adjustments in energy intake observed in females (Fig. 4D), in males is primarily driven by the increase in body weight over time (Fig. 4B), given the comparable calorie intake to chow-fed controls. Food intake percent changes (% Chng) were correlated with body weight variation (r=-0.7, p=0.001, Fig. 4J). We evaluated the hypothalamic expression of neuropeptides involved in the central regulation of energy balance as markers of anorexigenic and orexigenic signaling [23]. Their expression suggested sex-specific trends in animals fed with HFD relative to controls. While HFD females displayed higher levels of the anorexigenic transcript *Cart* compared to CD females and HFD males (Fig. 4F), the last exhibited lower expression of the orexigenic *Agrp* compared to CD males and HFD females (Fig. 4F). Consistently with its role, *Agrp* hypothalamic expression was correlated with % Chng in body weight and food intake (r=-0.45, p=0.04, and r=0.60, p=0.005, respectively) (Fig.4J). On the other hand, the expression of the associated satiety (*Pomc*) and hunger (*Npy*) neuropeptides did not differ between groups (Fig. 4F).

**Figure 4.**
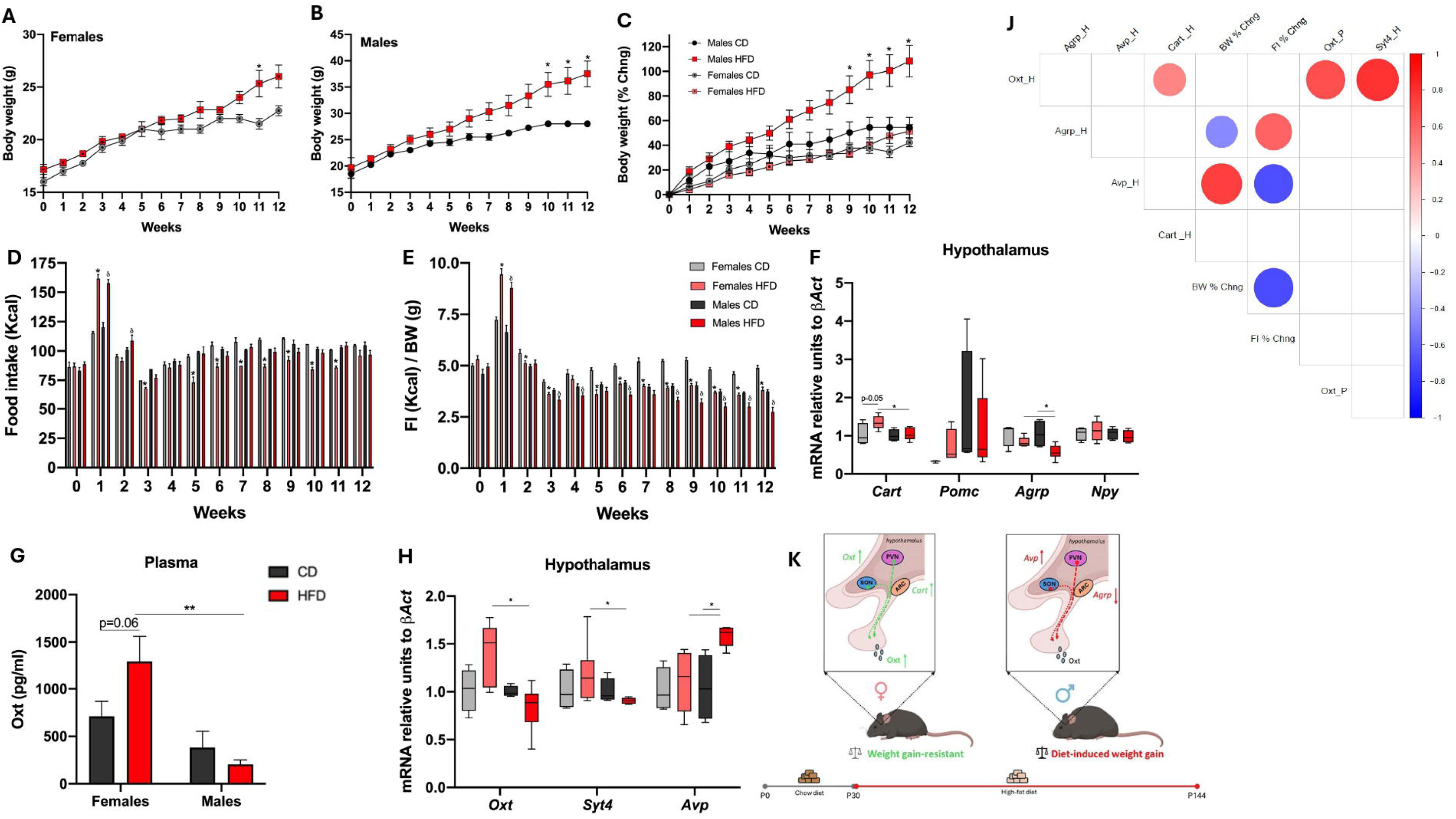
Sexual dimorphic Oxytocin and Vasopressin response to diet-induced obesity in mice. Body weight (BW) curves (g) over 12 weeks in female (**A**) and male (**B**) mice fed with chow diet (CD) or high-fat diet (HFD). Percentage change (% Chng) in BW (**C**). Food intake expressed in Kcal unadjusted (**D**) and adjusted for body weight **(E)** over 12 weeks of CD or HFD in males and females. qRT-PCR analysis of energy homeostasis regulatory genes: cocaine- and amphetamine-regulated transcript (*Cart*), agouti-related protein (*Agrp*), proopiomelanocortin (*Pomc*), and neuropeptide-y (*Npy*) (**F**) mRNA in the hypothalamus of CD and HFD males and females. Circulating Oxytocin (Oxt) plasma levels in male and female mice after 12 weeks of CD or HFD (**G**). qRT-PCR analysis of *Oxt*, Synaptotagmin-4 (*Syt4*), and Vasopressin (*Avp*) (**H**) in CD and HFD males and females. Correlogram illustrating correlations among circulating Oxt, hypothalamic gene expression (*Oxt*, *Avp*, *Cart*, *Agrp, Syt4*), and BW and FI %Chng (**J**). Conceptual scheme (created with BioRender) summarizing the sexually dimorphic neuroendocrine response to HFD (**K**). Comparisons between groups were performed with Student’s T-test. Data are expressed as mean ± SEM, and as mean and minimum and maximum level in F and H. *p<0.05; **p<0.01; ***p<0.001. *p<0.05 in female and δ<0.05 in male HFD vs CD in D-E. Pearson’s correlation coefficient (r) was calculated to assess the linear relationship between variables. PVN=paraventricular nucleus of the hypothalamus; SON=supraoptic nucleus of the hypothalamus; ARC= arcuate nucleus; p=postnatal day.

Pertaining to circulating Oxt levels, we observed overall higher plasma Oxt concentrations in these experiments compared to those performed at p21 and p60. This difference may be partly explained by the distinct dietary regimens (including differences in ingredients composition and macro- and micronutrient content between standard diet, CD, and HFD). Notably, mice at p21 were still breastfed, as this timepoint coincides with the weaning period. In addition, animals in the HFD experiment were analyzed at a different age (p120), which may have further contributed to the observed differences, as this later time point was necessary to allow sufficient duration for the dietary protocol to be performed.

HFD male mice displayed similar Oxt plasma concentrations compared to their normo-fed controls (Fig. 4H). On the other hand, in HFD-fed female mice a trend for higher Oxt plasma levels compared to their controls was detected (Fig. 4G, p=0.06). Overall, females with obesity displayed significantly higher circulating Oxt levels than males with obesity (Fig. 4G). These data were consistent with the hypothalamic *Oxt* expression which followed the same trend (Fig. 4H) and that was significantly correlated with Oxt plasma levels (r=0.68, p=0.002; Fig.4J) and *Cart* hypothalamic expression (r=0.48, p=0.04, Fig. 4J). Similarly to the data obtained in mice at weaning and in adulthood, *Syt4* hypothalamic expression paralleled that of *Oxt,* with HFD-fed females displaying higher levels than HFD-fed males (Fig.4H). Their levels were indeed significantly correlated (r=0.80, p<0.0001; Fig. 4J). The higher plasma Oxt levels observed in females with obesity, together with their relative resistance to weight gain, support the possibility that increased secretion of Oxt by the brain may contribute to reduced body mass gain in female mice exposed to HFD and may play a yet undefined role in the sex-dependent pathophysiology of obesity.

Lastly, *Avp* hypothalamic expression followed exactly the opposite trend as that of Oxt: HFD male mice exhibited higher levels compared to their CD-fed controls and to HFD females (Fig. 4H). Interestingly, *Avp* was strongly correlated with both % Chng in body weight (r=0.74, p<0.0001; Fig. 4J) and in food intake (r=-0.68, p=0.002; Fig. 4J). These data suggest that male and female mice may employ different neuroendocrine pathways in response to diet-induced obesity, with the first involving Avp and Agrp and the second Oxt and Cart, respectively (Fig. 4K).

## 4. Discussion

This study provides integrated data for understanding endogenous oxytocin biology across age, sex, and diet-induced obesity, combining precise nLC–MS/MS plasma measurements with hypothalamic gene-expression and morphometric analyses. We show that circulating Oxt levels rise from weaning to adulthood in both sexes and are mirrored by hypothalamic *Oxt* mRNA expression. Interestingly, Oxt neuron and IR-fiber densities follow sex-dependent developmental trajectories that diverge from plasma Oxt trends in males. Lastly, under HFD females exhibit higher circulating and hypothalamic *Oxt* levels compared to males and show relative resistance to diet-induced weight gain, pointing to a dimorphic Oxt-related response to obesity. These data provide a useful framework for interpreting previous studies and guiding future translational investigations.

Research on the Oxt system has increased steeply since the 1920s [1]. However, reliable quantification of plasma Oxt has been historically hindered by several well-recognized challenges. In the past years, the most common methods used to assess Oxt in plasma were commercially available immunoassays, which lead to contradictory results [7, 24]. For example, some studies report elevated concentrations, whereas others report no change [25] or even a reduction [25, 26] of Oxt levels in obesity (Table 1). Such inconsistencies have impeded progress in understanding Oxt physiology and its role in metabolic regulation. To overcome these limitations, we strictly controlled all confounders and employed a validated nLC–MS/MS, the gold standard technique for the detection of peptides with low molecular weight, including Oxt, in biological samples [19]. In humans, physiological Oxt levels measured by LC-MS are reported at 2.29±0.99 pg/mL (range: 1.05–3.67 pg/mL), whereas in rats, levels reach 1273±837 pg/mL (range: 398–3070 pg/mL) [27]. While Oxt levels have been extensively characterized in rats and humans through various methodologies [1, 27] there is still a notable lack of data for mice models specifically utilizing LC-MS. In the present study conducted in mice, plasma Oxt concentrations showed a wide range, from 15 to 2192 pg/mL, considering both physiologic and obesity conditions. To our knowledge, the present study represents a preliminary effort to quantify murine Oxt plasma levels via LC-MS/MS.

A key observation is that, in both sexes, Oxt plasma levels and hypothalamic expression similarly rise from weaning to adulthood, indicating coordinated transcriptional and release upregulation. However, Oxt neuron and IR-fiber densities show sexually dimorphic and age-dependent patterns: in adulthood, females exhibit increased Oxt neuron density and denser hypothalamic projections, whereas males exhibit reductions, in both PVN and SON. These findings integrate with existing reports showing that Oxt neuron number increases from p1 to p21 in both sexes [28], and that adult females display a higher number of Oxt neurons than males [29]. Sex dimorphism in fibers distribution and prevalence across different brain regions has been controversial: one study reported higher number of fibers projecting towards mesolimbic regions in female adult mice [29] while genetic labeling reported similar projections between sexes in 4 months old mice across different brain areas [30]. Notably, our study established higher Oxt-IR fiber density in females compared to males at p21 and p60 in both, PVN and SON, location projecting to the median eminence [22]. The sexually dimorphic trajectories in Oxt neuron and IR-fiber densities observed between p21 and p60 likely reflect the influence of sex hormones on the Oxt system, which becomes significant following puberty. This period, which in mice spans from _∼_p26 to _∼_p40, is characterized by rising levels of sex steroids and the achievement of sexual and reproductive maturity. Importantly, while testosterone administration in gonadectomized mice was shown to reduce the number of neurons in PVN and SON [31], estrogen administration promoted Oxt mRNA expression and release [17].

Interestingly, our concerted assessment of Avp reveals distinct developmental and regulatory trajectories for the two closely related nonapeptide’s systems. Unlike Oxt, where plasma levels and hypothalamic expression increase coordinately with age in both sexes, Avp exhibits a sexually dimorphic and divergent pattern: plasma Avp rises in adult females but decreases in males, with opposing trends in hypothalamic expression. This uncoupling highlights a distinct regulatory logic between the two nonapeptides, already pointed by others [32], and that deserve further investigation.

Sexual differences in the neuroendocrine response to HFD-induced obesity emerged as a major finding. During the first week of HFD exposure, both sexes showed increased caloric intake, likely due to the higher palatability of the diet. Thereafter, females reduced their caloric intake relative to controls and displayed a similar body weight, whereas males did not exhibit a similar adjustment in food intake and instead showed greater weight gain. These results indicate that macronutrient composition, rather than absolute caloric intake, may contribute to the greater body weight gain observed in males, as caloric intake was comparable between CD- and HFD-fed males for most of the study period. Importantly, this data also suggests a more effective compensatory response to HFD in females. The higher metabolic resilience of C57BL6/J females in response to HFD is established in the literature and consistent with our data [33]. Female mice exposed to such dietary protocol experience a slower increase in body weight, elevated energy expenditure, with no variation in locomotor activity, and better glucose control compared to males [34]. Importantly, HFD-exposed females displayed significantly higher plasma Oxt levels than males, paralleled by increased hypothalamic expression of *Oxt* and of the anorexigenic gene *Cart*. These results suggest that higher circulating Oxt levels in females with obesity may contribute to their relative metabolic resilience compared to males. Beyond its role as a central satiety regulator, Oxt is known to promote lipolysis, thermogenesis, and energy expenditure through a peripheral action [5, 6], processes that could be activated in female mice with obesity who display elevated Oxt levels. Conversely, males, who were more susceptible to diet-induced weight gain, did not show changes in plasma Oxt but exhibited a marked increase in hypothalamic *Avp* expression and a reduction in the orexigenic regulator *Agrp*. Similarly to Oxt, Avp was demonstrated to reduce food intake, to display a broad metabolic action, and based on our data it is upregulated to respond to diet-induced obesity in males [35]. Based on our data it is hence possible that females preferentially rely on Oxt and Cart pathways, whereas males may employ Avp and Agrp pathways under metabolic stress, highlighting a divergence that may contribute to the greater male susceptibility to obesity and metabolic complications, a sex-specific response that warrants further investigation (Fig. 4K).

Although our study has some limitations, most notably the lack of mechanistic experiments needed to fully delineate the sexually dimorphic regulation of Oxt and Avp, as well as the absence of data on energy expenditure, our results provide clear avenues for future research. Further studies assessing energy expenditure in the HFD model and measuring circulating Avp levels will help refine the regulatory framework we propose. Nonetheless, through a rigorous and integrated methodological approach, our study helps resolve long-standing inconsistencies in mice Oxt biology and establishes a reference point for future mechanistic and translational investigations, highlighting the importance for future studies to explicitly control for age and dietary composition when evaluating circulating Oxt levels, as these factors appear to play a critical role in determining plasma concentration.

## Supporting information

Supplementary figure 1, Methodological validation of neuropeptide quantification in plasma and oxytocin antibody specificity.

## Author Contribution

G.C., C.G. and A.G.: Conceptualization; Writing-Original Draft; G.C., C.G.: Data Curation and Formal Analysis; G.C and A.G. Funding Acquisition; G.C., C.G., S.M., S.D.B., I.S., J.P., E.S. G.G.: Methodology, Investigation, Data Curation. G.Cr., A.G.: Supervision; G.C., C.G., S.M., S.D.B., I.S., J.P., E.S., G.Cr., A.G.: Writing-Review & Editing.

