## Supplementary figures and images for "Plasma oxytocin measured by LC–MS/MS varies with life stage, sex, and obesity in mice"

### Supplementary figure 1, Methodological validation of neuropeptide quantification in plasma and oxytocin antibody specificity.

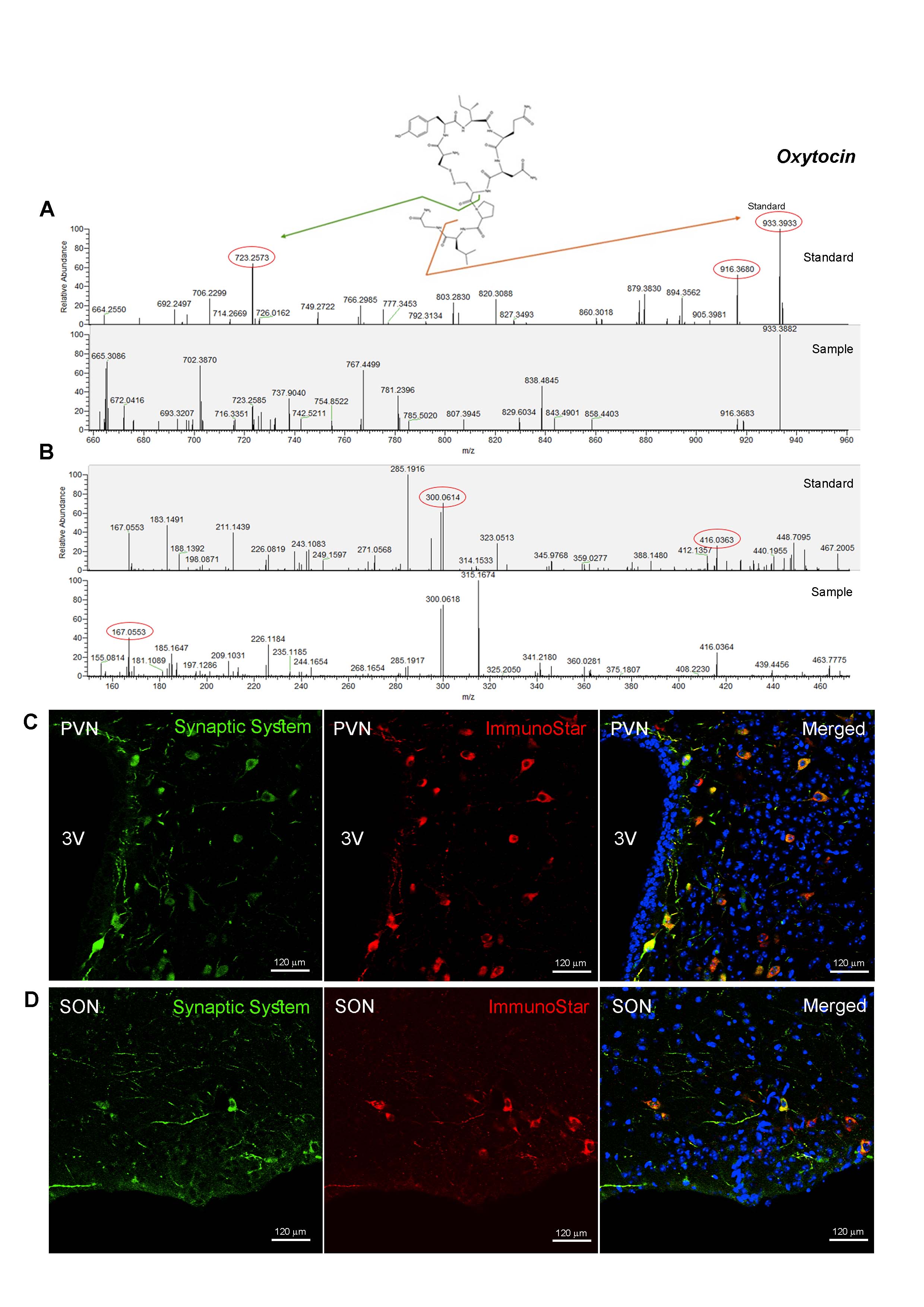
