## Supplementary table 1,2,3a,3b + supplementary figure 1 legend for "Plasma oxytocin measured by LC–MS/MS varies with life stage, sex, and obesity in mice"

**Supplementary Materials**

***Supplementary Table 1***

| PRIMERS | |
|---|---|
| **Target gene** | **Sequence** |
| *Oxt* | F: CTTACTGGCTCTGACCTCGG<br>R: CACTTGCGCATATCCAGGTC |
| *Syt4* | F: ACCCCAAAGCTCTTTACGGA<br>R: GAGTGTCCCCAGCTTCTCTT |
| *Avp* | F: ACTTCCAGAACTGCCCAAGA<br>R: GGTAGTTCTCCTCCTGGCAG |
| *Cart* | F: TACTCTGCCGTGGATGATGCGT<br>R: TCGGAATGCGTTTACTCTTGAGC |
| *Agrp* | F: ATGCTGACTGCAATGTTGCTG<br>R: CAGACTTAGACCTGGGAACTCT |
| *Pomc* | F: CCATAGATGTGTGGAGCTGGTG<br>R: CATCTCCGTTGCCAGGAAACAC |
| *Npy* | F: CAGAAAACGCCCCCAGAA<br>R: AAAAGTCGGGAGAACAAGTTT |
| *β-actin* | F: GTGGGAATGGGTCAGAAGGA<br>R: CTTCTCCATGTCGTCCCAGT |

***Supplementary Table 2***

| PRIMARY ANTIBODIES | | | |
|---|---|---|---|
| **Antibody** | **Source** | **Dilution** | **Brand** |
| Oxytocin | Polyclonal guinea pig | 1:8000 (IDAB)<br>1:7000 (IF) | Synaptic System (#408 004) |
| Oxytocin | Polyclonal rabbit | 1:16000 (IF) | ImmunoStar (#20068) |
| **SECONDARY ANTIBODIES** | | | |
| **Target and conjugate** | **Source** | **Dilution** | **Brand** |
| Anti-Guinea Pig IgG biotinylated | Goat | 1:200 | Vector Laboratories (#BA-7000-1.5) |
| Anti-guinea pig Alexa Fluor 488 | Donkey | 1:200 | Jackson ImmunoResearch (#706-546-148) |
| Anti-rabbit Alexa Fluor | Donkey | 1:400 | Invitrogen (#A31572) |

***Supplementary Table 3a*** *– Trueness and precision*

| Oxytocin | | | | | | | |
|---|---|---|---|---|---|---|---|
| Validation level (ng/L) | Day | Apparent recovery (%) | Intraday CV (%) | Inter-day CV (%) | Pooled apparent recovery (%) | $CV_{r,\ pooled}$ (%) | $CV_{R,\ pooled}$ (%) |
| 2.5 | 1 | 83 | 4.1 | 13.0 | 100 | 7.0 | 10.2 |
| | 2 | 95 | 16.3 | | | | |
| | 3 | 95 | 12.0 | | | | |
| 10 | 1 | 104 | 6.0 | 8.4 | | | |
| | 2 | 103 | 5.4 | | | | |
| | 3 | 89 | 2.1 | | | | |
| 50 | 1 | 110 | 5.5 | 8.5 | | | |
| | 2 | 95 | 7.0 | | | | |
| | 3 | 108 | 5.9 | | | | |
| 100 | 1 | 112 | 5.9 | 8.8 | | | |
| | 2 | 94 | 5.9 | | | | |
| | 3 | 106 | 3.1 | | | | |
| 500 | 1 | 107 | 8.4 | 8.8 | | | |
| | 2 | 110 | 6.3 | | | | |
| | 3 | 95 | 4.5 | | | | |
| 1000 | 1 | 113 | 5.9 | 8.6 | | | |
| | 2 | 107 | 4.8 | | | | |
| | 3 | 95 | 2.7 | | | | |

*__Supplementary Table 3b__ – Trueness and precision*

| Vasopressin | | | | | | | |
|---|---|---|---|---|---|---|---|
| Validation level (ng/L) | Day | Apparent recovery (%) | Intraday CV (%) | Inter-day CV (%) | Pooled apparent recovery (%) | $CV_{r, pooled}$ (%) | $CV_{R, pooled}$ (%) |
| 25 | 1 | 117 | 15.5 | 13.4 | 103 | 8.3 | 9.8 |
| | 2 | 116 | 8.0 | | | | |
| | 3 | 108 | 11.8 | | | | |
| 50 | 1 | 95 | 14.3 | 12.2 | | | |
| | 2 | 99 | 12.0 | | | | |
| | 3 | 111 | 6.0 | | | | |
| 100 | 1 | 96 | 3.1 | 5.1 | | | |
| | 2 | 97 | 5.7 | | | | |
| | 3 | 102 | 4.9 | | | | |
| 500 | 1 | 106 | 4.5 | 4.9 | | | |
| | 2 | 98 | 3.1 | | | | |
| | 3 | 101 | 36.5 | | | | |
| 1000 | 1 | 106 | 4.9 | 6.2 | | | |
| | 2 | 98 | 1.5 | | | | |
| | 3 | 97 | 7.8 | | | | |

***Supplementary Figure 1***

**Figure S1.** *Methodological validation of neuropeptide quantification in plasma and oxytocin antibody specificity*. Detailed MS/MS spectra of Oxt in both standard solution and biological samples with presumptive annotations for the characteristic fragments at high **(A)** and low **(B)** mass. For the *m/z* 300.0614, it was only possible to assign a theoretical formula corresponding to $[C_{14}H_{11}O_5N_3]^+$ (Rings and Double Bonds: 11.5), resulting from a rearrangement of the cyclic moiety of oxytocin that includes an aromatic ring. **(C)** Double-labeled confocal microscopy for Oxt in the PVN and the SON showing the colocalization between two different primary antibodies directed against Oxt from Synaptic System and ImmunoStar **(B)**. Standard (oxytocin-[leucine-5,5,5-d3, glycine-2,2-d2] trifluoroacetate salt); PVN=paraventricular nucleus of the hypothalamus; SON=supraoptic nucleus of the hypothalamus; 3V=third ventricle.
